# Spatial modeling of Cultural Ecosystem Services from social media data: Systematic review of operability, opportunities, limitations and ways forward

**DOI:** 10.1101/2025.07.16.665139

**Authors:** Carlos Javier Navarro, Akram Elghouat, Marta Rueda-Pasadas, Jose Carlos Pérez-Girón, Ricardo Moreno-Llorca, Ana Sofia Vaz, Javier Martínez-López, Nuria Pistón, Domingo Alcaraz-Segura

## Abstract

1. This systematic review explores the use of social media data to spatially model Cultural Ecosystem Services (CES), which are non-material benefits provided by ecosystems that support human well-being. Based on a search of 510 scientific articles in Web of Science and Scopus, we carefully selected and analyzed those that focused on spatial modeling of CES using social media data. We aimed to (a) identify the diversity of CES assessed, (b) analyse the social media platforms used as data sources, (c) evaluate the modelling frameworks employed, and (d) summarise the predictor variables included in these models.
2. We found that the most studied CES were those related to physical and psychological experiences (62%; especially recreation) and the main predictor variables were the presence of natural elements (52%), land use and land cover maps, and topographic variables, often weighted by applying distance-based metrics (24%). Twenty-four social media data sources were identified but Flickr was by far the most widely used one (40%). MaxEnt (37%) and Random Forest (16%) were the most commonly used modeling tools. The most commonly used metrics to assess model performance were AUC-ROC, AIC, and R2 values.
3. While the use of social media offers an opportunity to study CES and provide cost-effective and scalable insights, this article discusses some limitations and considerations raised from the literature review to be taken into account when using this type of data. These include the quality and representativeness of social media data, the importance of a clear definition of CES, a proper labeling of social media data, and an appropriate selection of spatial modeling techniques.
4. Future research should address these limitations and considerations by integrating different data sources and refining methodologies to improve the accuracy and applicability of CES models. This review provides a comprehensive overview of current practices and highlights areas for further investigation in the spatial modeling of CES using social media data.

## 1. Introduction

### Definition of CES

Cultural Ecosystem Services (hereafter called CES), are defined as non-material outputs of ecosystems that affect the physical and mental states of people (MEA 2005; Haines-Young and Potschin 2018). CES include benefits, such as improved physical and psychological well-being, opportunities for recreation and social interaction, intellectual and spiritual benefits, connections to socio-cultural heritage, and biodiversity appreciation, among others (Nowak-Olejnik et al. 2022). CES have been subject to different theoretical and terminological interpretations, depending on the framework adopted (De Groot et al. 2018). Similar terms include non-material nature’s contributions to people which represent the impact of nature on subjective and psychological aspects that contribute to an individual’s overall quality of life (Díaz et al. 2018). Since the emergence of the concept of ecosystem services, much scientific research has been published with particular emphasis on provisioning and regulating services (Kremen and Ostfeld 2005; Schaich et al. 2010; Hernández-Morcillo et al. 2013). The study of CES has historically been less prevalent, although there is recently a growing interest in its valuation (IPBES 2018). One of the Sustainable Development Goals (UN, 2015), and of particular relevance to the European Union in particular, is the conservation of nature and the CES. While significant progress has been made in incorporating provisioning and regulating services, especially in the management of protected areas (e.g. Castro et al. 2015), CES still remains significantly underrepresented (MINAM, 2014; Costanza et al. 2017; Lin et al. 2024). Protected areas play a pivotal role, serving not only to safeguard flora and fauna, but also to provide multiple CES (Zorrilla-Miras et al. 2014; Cheng 2023), within their boundaries as well as in the surrounding areas (Palomo et al. 2013; Zhao et al. 2023). In this regard, integrating ecosystem services evaluation into land use planning and management remains a challenge, mainly due to the lack of policies mandating their inclusion and the limited availability of accurate, standardised maps at scales suitable for local decision-making (Albert et al. 2014).

### Quantification of CES

Different methodologies have been developed to quantify what aspects of nature people value, enjoy, and are more interested in. These methodologies range from surveys, participatory mapping, analysis of tourism records or government statistics (Martín-López et al. 2012; García-Nieto et al. 2013; Castro et al. 2014; Langemeyer et al. 2023; Cheng 2023). Within these methodologies, the use of social media for the quantification of CES stands out. This approach has a series of advantages such as being relatively easy to obtain, representing a direct input from social media users, and being able to cover large spatial and temporal scales (Sonter et al. 2016). A variety of analytical approaches may be employed in the examination of this type of data, including the automated analysis of image content, semantic analysis of social media posts or reviews, and the analysis of the distribution of geolocated images, posts, routes, and points (Manley et al. 2022). However, like any other technique, quantifying CES from social media also has some limitations, such as the lack of precision in the location of photos, limitations to access the information, the challenge of filtering relevant content from a large amount of irrelevant information, among others (Otero-Rozas et al. 2018; Ghermandi and Sinclair 2019). In addition, content biases arising from the type of user (e.g. age range, socio-economic characteristics) and the nature of information shared on social media need to be addressed (Hargittai 2020). And spatial biases, such as overrepresentation of easily accessible locations or gaps created by lack of internet access (Pick et al. 2024).

### How to map and model CES

One straightforward way to use social media data is to model the occurrence and abundance of geolocated information, when available, by means of species distribution (SDM) and abundance (SAD) models (Cheng 2023). Although this type of modeling was originally conceived to quantify the probabilities of occurrence of animal or plant species, known as niche or habitat suitability, it has been successfully used in other contexts (Sillero et al. 2021), such as to predict fire occurrence (Batllori et al. 2013), to define protected areas (Wang et al. 2016), or to map Ecosystem services (Richards and Friess 2015; Lavorel et al. 2017) and more specifically CES (Clemente et al. 2019;Arlsan and Örücü 2021; Ma and Yang, 2025). These models are mathematical representations of the CES’s niche that integrate the drivers that characterize the environmental suitability of individual CES and that project it onto the geographic space (Guisan et al. 2017; Sillero et al. 2021). This type of modelling usually incorporates a wide range of drivers or predictor variables that define the composition, structure and functioning of the social-ecological system including climatic, topographic, accessibility, diversity of habitats and land use, or primary production among others (Vaz et al. 2020). Initially, spatial modelling of CES occurrence did not distinguish between different types of CES but analysed all of them as a whole (Posner et al. 2016). However, recent evidence suggests that specific CES (i.e. narrower subtypes of CES) show stronger and more robust correlations with environmental variables than broader classes (Hale et al. 2019; Lingua et al. 2022), as is often the case when modelling the habitat of the different plant or animal species within a genus or family (Guisan and Zimmermann 2000).

### Objectives

In this article, we present a systematic literature review of studies dealing with spatial modeling of CES from social media data, with the aim of compiling and analyzing: a) the variety of CES assessed, b) the main sources of social media data, c) the modeling framework, and d) the predictors or explanatory variables used.

## 2. Materials and methods

### 2.1 Literature selection criteria

The present literature review was conducted in accordance with the PRISMA methodology (Page et al. 2021), a structured and transparent approach for systematic reviews. This methodology encompasses the comprehensive identification of pertinent studies, the removal of duplicates, the evaluation of eligibility based on predefined criteria, and the inclusion of selected studies. To identify relevant scientific articles dealing with spatial modeling of CES, we focused exclusively on English-language scientific articles (excluding literature reviews) published up to 31/December/2024 in the Web of Science (WOS) and Scopus databases. We searched for studies containing at least one CES-related term (cultural ecosystem service*, ecosystem service*, nature* contributions to people, nature’s contribution* to people, nature* benefit to people, nature*based solution*, human*nature relationship*, human*nature connect*, nature gift, material contribution*, non*material contribution*, environment* service*, ecologic* service*, ecosystem function*, environment* function*, ecologic* function*) co-occurring with at least one Social media term (social media, Twitter, Flickr, Instagram, Facebook, Social networks) in the title, summary, or authors’ keywords.

This search returned 314 scientific articles in WOS and 487 in Scopus. After removing duplicates, a total of 510 unique articles remained for analysis. After analyzing the titles and abstracts and excluding articles that did not use social media to quantify CES, 123 articles were selected for eligibility. Finally, upon detailed reading and analysis, we further excluded 65 articles that did not focus on CES modeling. Therefore, this processing resulted in a final selection of 58 articles dealing with spatial modeling of CES. For a better understanding of the peer-review process, the workflow is exemplified in Figure 1.

**Figure 1.**
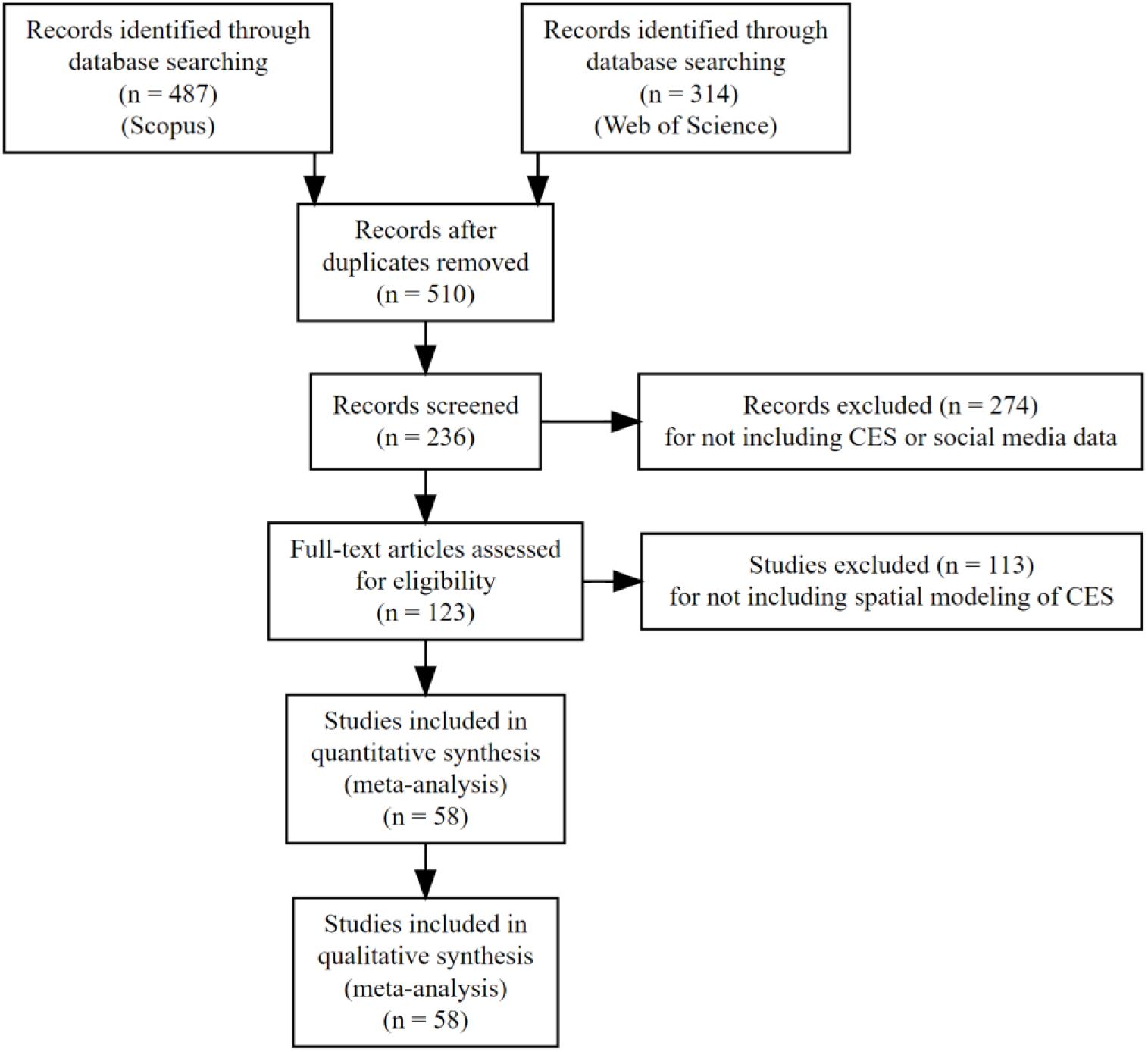
Flow chart of peer-review procedure based on PRISMA method (adapted from Page et al. 2021).

### 2.2 Metadata coding

From each article, we extracted the type of CES addressed, the social media data source, the explanatory variables used, the modeling frameworks, and other complementary information as described below (Table S1 in Supplementary material).

#### 2.2.1 Type of CES modeled

First, we identified and read each definition of CES in each article. When identifying an inconsistency between the definition of CES and the category assigned to that CES by the authors of the articles, we established a semantic consensus based on the precise type of CES that was addressed in the article. To this end, we adhered to the categories proposed by Díaz et al. (2018) in order to standardize the categorisation of CES across published studies. For example, a CES that has been designated by authors as a single category but, in fact, in the definition given by the authors encompassed two or more different CES categories as outlined by Díaz et al. (2018). Clemente (2019), for example, refers to the scientific and educational CES, which for Díaz are two different CES: “research” and “environmental education”. In such cases, different CES by its definition were recorded separately to be consistent with the standardization protocol adopted (see Table S3 for a detailed description). In addition, for each CES analysed, we noted whether it focused on supply, demand or both. In cases where this was not explicitly stated, we have recorded it as undefined.

#### 2.2.2 Social media data source

To gain an overview of the platforms that are used as sources of social media data, we identified the following information for each article: the platform where data were obtained (e.g. Flickr, X-Twitter, Instagram), the type of data (geolocated photos, geolocated text, location, etc.) and the temporal coverage.

#### 2.2.3 Explanatory variables

To obtain information on the explanatory variables used, we extracted the predictors used from each article. To this end, we standardized the variables: if the variables were exactly the same but named differently (e.g. altitude = height), we standardized their names for consistency. We then identified the aggregation methods used to calculate each variable, as it is common for a variable to be calculated using different aggregation methods (e.g. distance to water bodies, density of water bodies, proportion of water bodies). Once standardised, we grouped the variables into three-level hierarchical groups, based on the groupings proposed above in the literature and our own criteria (Table S1). In cases where several CES were evaluated, the variables used (if they used the same explanatory variables) were counted as many times as CES.

#### 2.2.4 Spatial Modelling

To compile information about the models, we extracted the modelling approach used in each article. In the articles that used more than one model type, we also indicated the model that performed the best.

#### 2.2.5 Model evaluation metrics

To assess the performance of the models used in the reviewed literature, we also considered evaluation metrics commonly used in spatial modelling. These metrics allow us to compare the ability of models to capture the distribution of CES based on the social media data used.

#### 2.2.5 Spatial cover and ecosystem types

We also extracted information such as spatial extent, categorized as local (area <100 km^2^), regional (area >100 km^2^) or national, continental and global scale. Additionally, the countries where the studies were conducted were recorded. The type of ecosystem in which the studies were conducted were recorded following The Economics of Ecosystems and Biodiversity (TEEB) classification for biomes and ecosystems, which comprises 15 biomes and 84 ecosystem types (Kumar 2012; Brander et al. 2024). If the study was conducted across multiple ecosystems, the three predominant ecosystem types were included. All information extracted from the articles is presented in Table S1 in the supplementary material.

## 3. Results

From the 58 sets of studies, we found that research in this topic is mainly concentrated in China and the United States, with an increasing number of articles since 2015 (Figure 2). Most of the studies were regional (60%), followed by local (28%) and national ones (12%).

**Figure 2.**
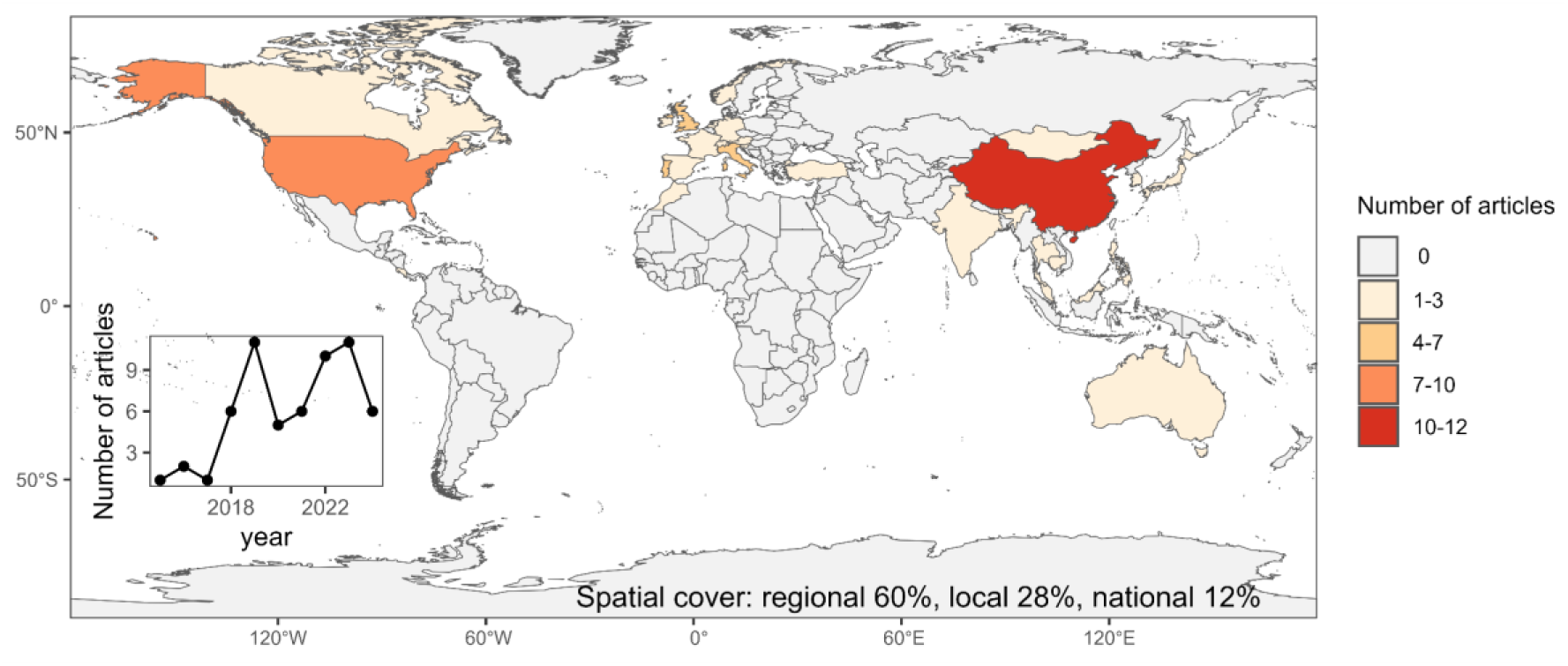
Distribution per country and across years of the number of articles indexed in Scopus and Web Of Science that dealt with the spatial modeling of cultural ecosystem services (CES) using social media data.

### 3.1 Type of CES modeled

Of the studies analyzed, 21% addressed both supply and demand simultaneously, 24% focused exclusively on supply, 24% addressed only demand and 31% lacked a clearly defined focus. In terms of CES categories, studies focusing on physical and psychological experiences were the most represented (61%), with a focus on recreational activities, enjoyment of the landscape and enjoyment of fauna and flora. Secondly, 22% of the studies focused on the category of supporting identities, related to cultural heritage and spiritual significance. Furthermore, CES related to learning and inspiration, such as environmental education and artistic inspiration, accounted for 23% of the revised articles (Figure 3).

**Figure 3.**
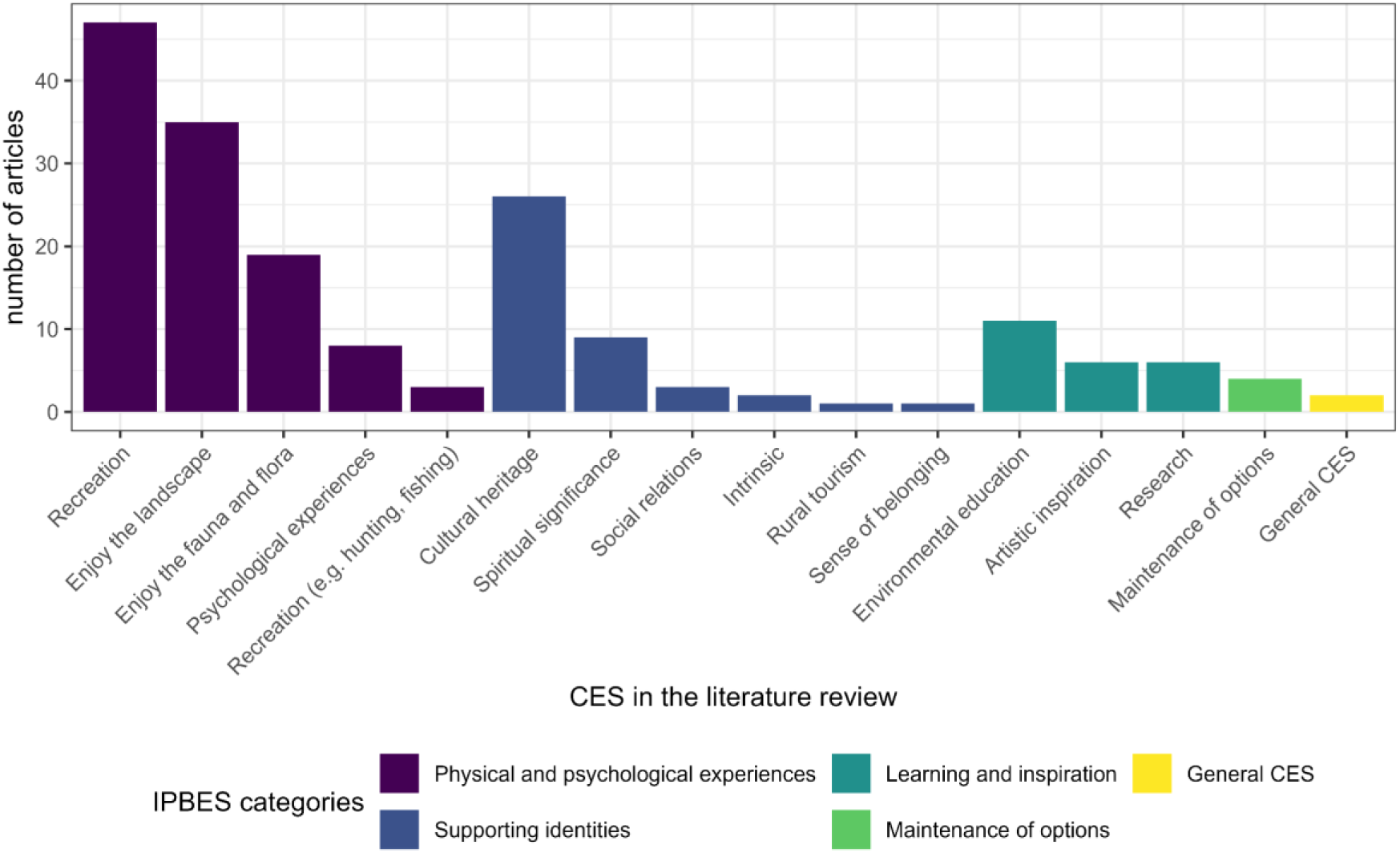
Type of cultural ecosystem services (CES) evaluated in the analyzed articles according to the IPBES (Intergovernmental Science-Policy Platform on Biodiversity and Ecosystem Services) categorization and a more detailed description of Nature’s Contributions to People (NCP’s) within each IPBES (2018) category (X axis).

We identified a total of 184 unique CES, classified into 15 broad types of CES according to Diaz et al. 2018 using the definitions provided by the authors. Of the total of CES identified, approximately 30% did not match the CES definitions assigned to it by authors and had to be recategorised (Table 1). For details on how we categorized the CES, see Table S2.

**Table 1.**
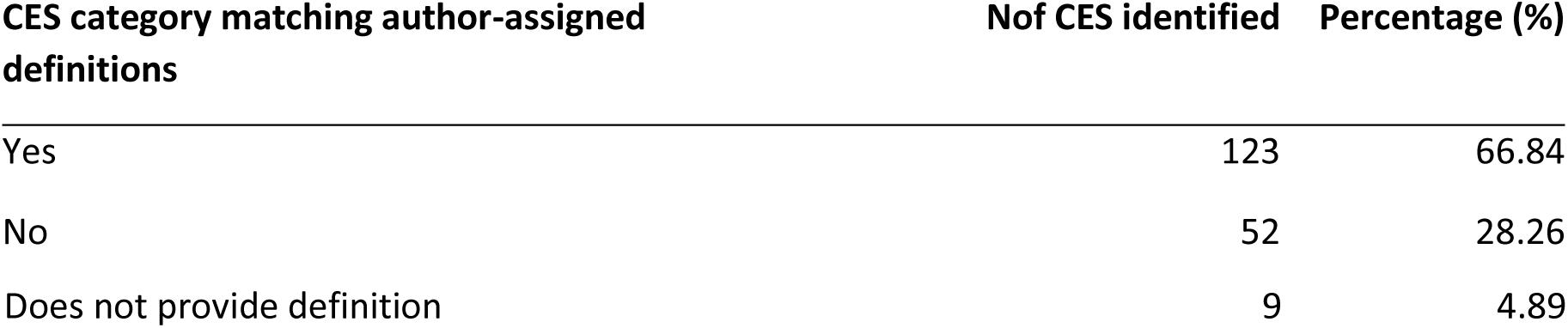
Number of cultural ecosystem services (CES) (in many cases more than one CES per article) for which there was consensus between the authors’ definition and the CES category originally assigned (semantic consensus).

### 3.2 Social media data source

We found that Flickr was the most frequently used data source in the articles we analyzed, accounting for 40% of the total (Figure 4). The predominant type of data was geolocated photos, representing 63% of the total, although in some cases text, geolocation coordinates, sign-in data, etc. were also used.

**Figure 4.**
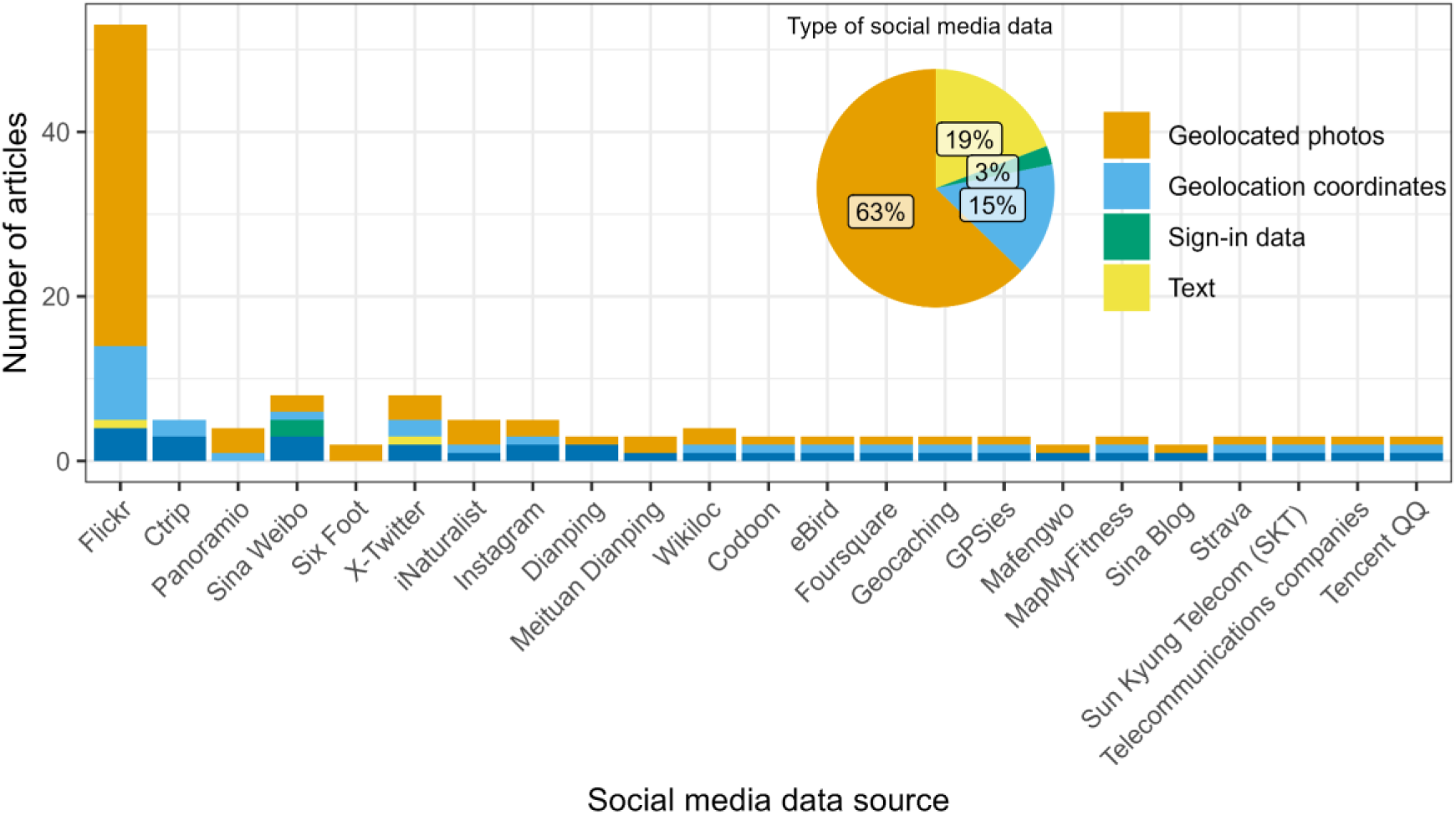
Distribution of social media data used by source and type of data collected. The bar chart shows the frequency of data sources. The pie chart shows the relative percentage of each type of data.

### 3.3 Explanatory variables

Explanatory variables were most frequently classified in the category of natural elements (52%; level 1), including rivers, dams and mountains (level 2), followed by the category of land use and the levels of land cover and accessibility (Figure 5). We identified a total of 362 variables, once standardized, with the most common being roads (112), elevation (72) and land use/land cover (72), furthermore, these variables are present as predictor variables for all CES groups (Table 2). See Table 2 for details of the remaining CES, and Table S2 for an overview of the original variables of the articles. The most common aggregation methods from the original layers were distance (24%) and counts (12%) (Figure S1).

**Figure 5.**
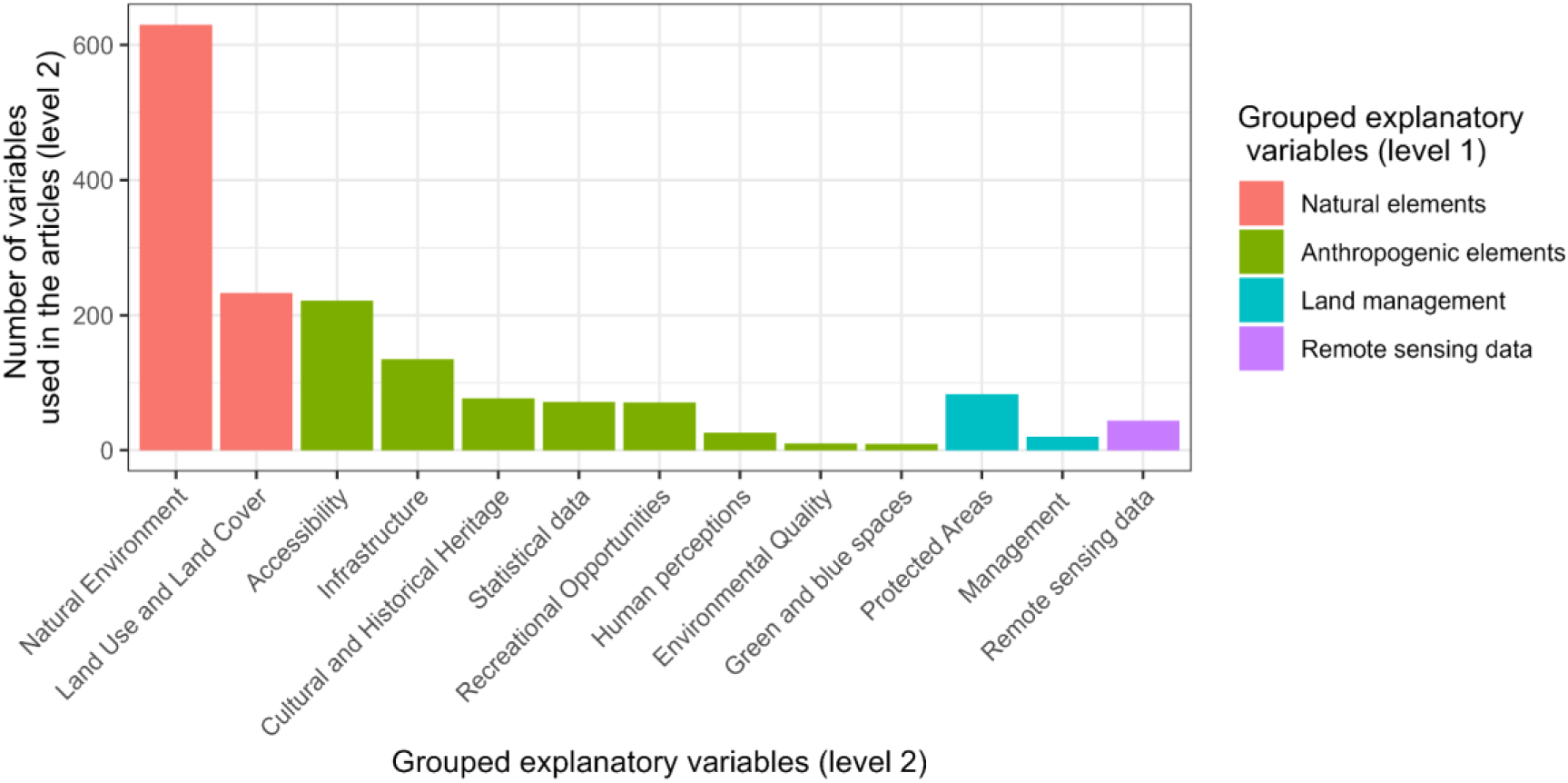
Variables used as predictors in cultural ecosystem services (CES) modelling. The most common variables are shown by type of CES analysed. The variables have been grouped hierarchically for ease of visualization. The x-axis shows the variables grouped at their level 2, while the labels and colors refer to level 1. The original variables can be found in Table XX in the supplementary material.

**Table 2.**
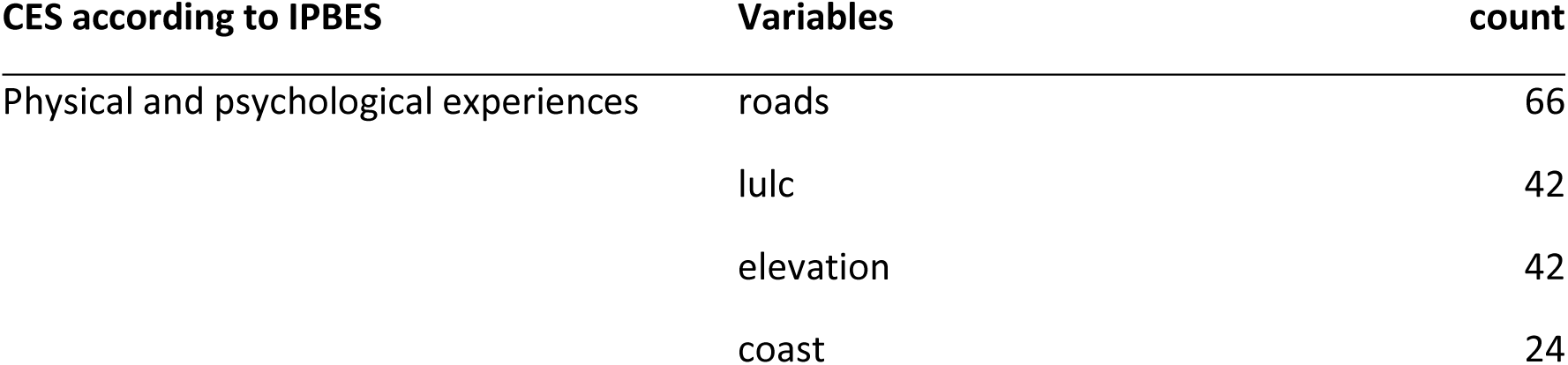
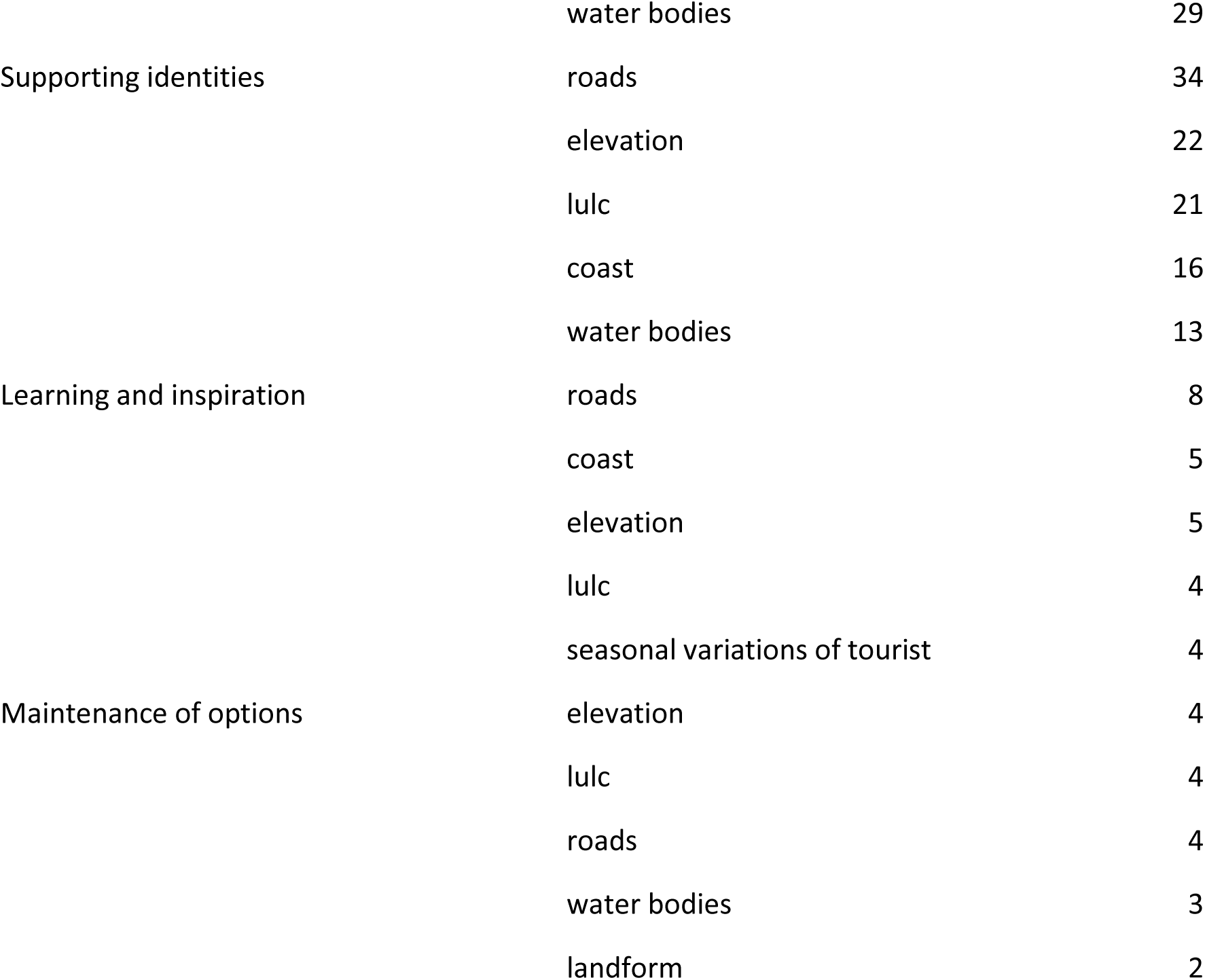
Variables most commonly used as predictors in spatial modeling of cultural ecosystem services (CES, grouped by type of CES according to IPBES. The five most commonly used variables in CES modelling are shown.

### 3.4 Spatial Model

Regarding the models used in the spatial modeling of CES, the predominance of MaxEnt (used in 37% of articles) stands out, followed by Random Forest (16%) and Simple Linear Regressions (13%). Figure 6 shows the number of times each model was used, where only one model was used (in green) and also in the cases where multiple models were used, one was selected as the best model (orange). In general, the assembly of different types of models is not usually used, although we found an article where they do work with this approach (Fox et. al 2021).

**Figure 6.**
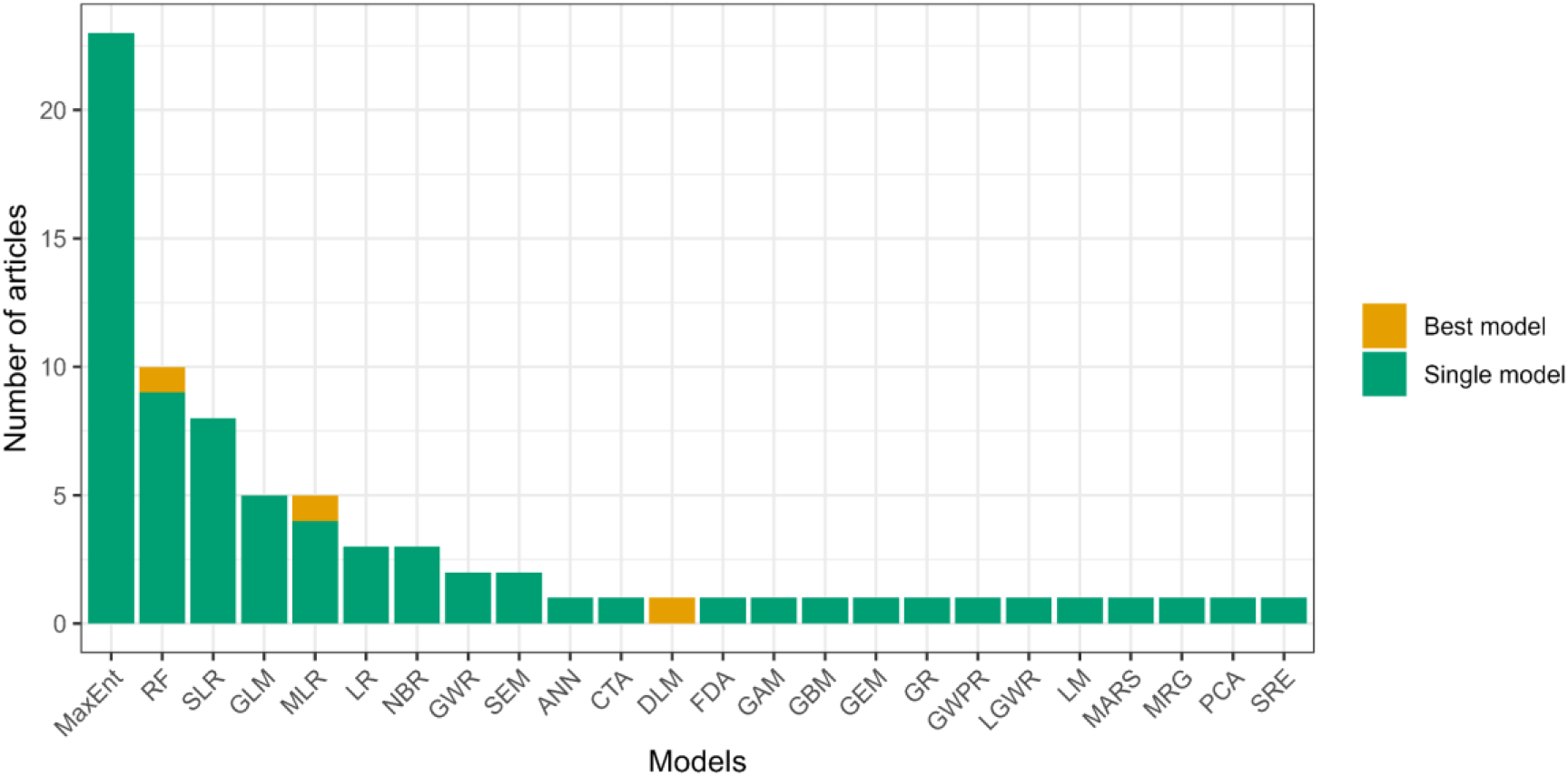
Models used in the analyzed articles. The models that were present in articles that evaluated multiple models and where they were the best are in orange, the rest of the models in green. ANN (Artificial Neural Network), CTA (Classification Tree Analysis), DLM (Dual Logarithmic Model), FDA (Flexible Discriminant Analysis), GAM (Generalized Additive Model), GBM (Generalized Boosting Model), GEM (Global Ecological Model), GLM (Generalized Linear Model), GR (Global Regression), GWR (Geographic Weighted Regression), LGWR (Logistic Geographic Weighted Regression), LM (Logarithmic Model), LR (Logistic Regression), MARS (Multiple Adaptive Regression Splines), MaxEnt (Maximum entropy), MLR (Multiple Linear Regression), NBR (Negative Binomial Regression), OLS (Ordinary Least Squares), RF (Random Forest), SEM (Structural Equations Models), SRE (Surface Range Envelope). Articles where a single model was used are in green, while those models where a multi-model approach was used but only one was chosen as the best model is in orange.

The most commonly used evaluation metrics were the area under the curve (AUC-ROC) (29.85 %), the coefficient of determination (R² and pseudo R²) (25.37 %) and the Akaike information criterion (AIC) (16%). The average values of the two most commonly used evaluation metrics indicate that the AUC typically used is around 0.85, while the R² value is 0.22, although with considerable variability, as shown in Figure 7.

**Figure 7.**
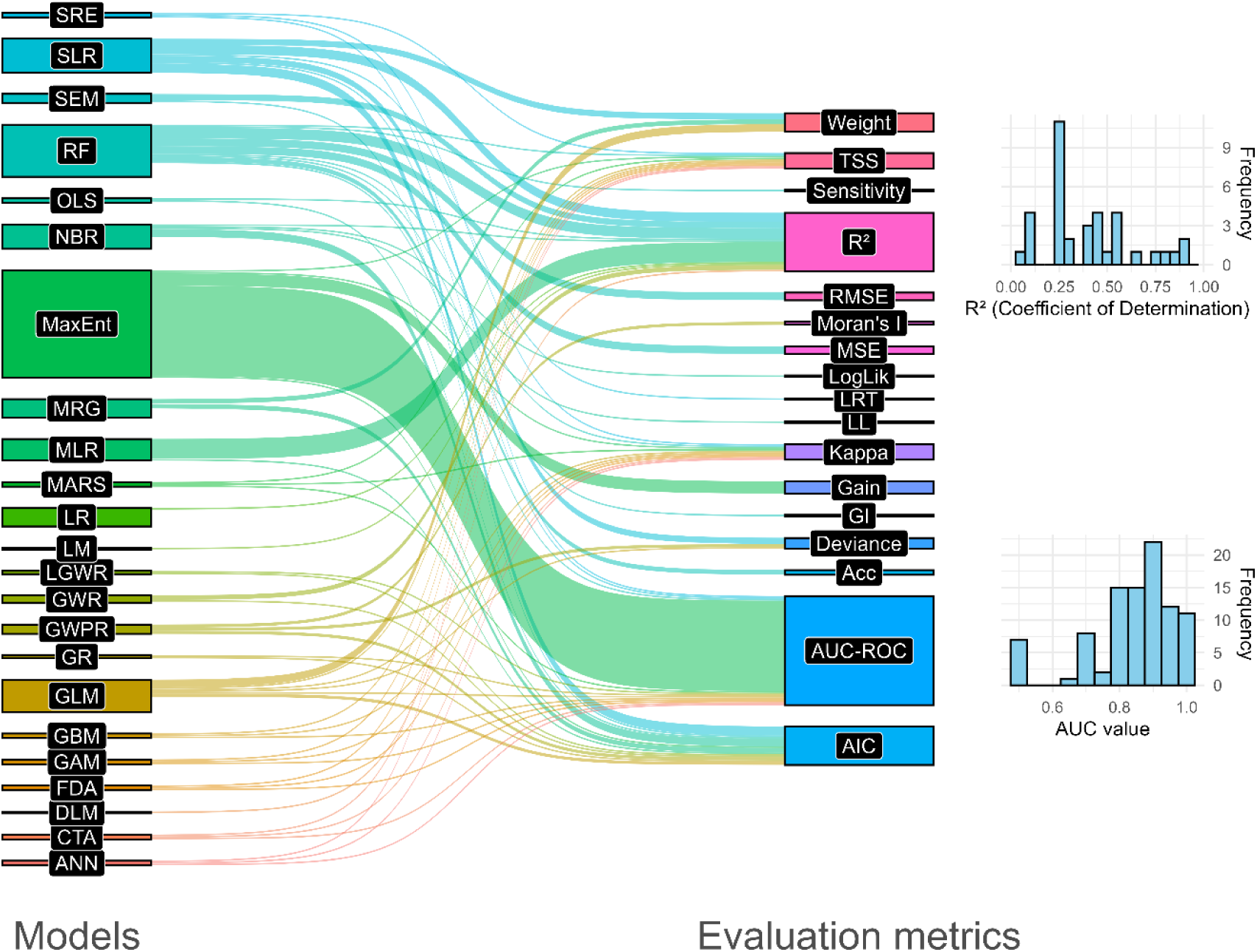
Relationship between different models and evaluation metrics used for their comparison in the analyzed articles. See Figure 6 and its caption for model descriptions. Evaluation metrics: AIC (Akaike Information Criterion), AUC-ROC (Area Under the Receiver Operating Characteristic Curve), Acc (Accuracy), Gain (Gain), GTI (Goodness of Fit Index), LRT (Likelihood Ratio), LogLik (Log-Likelihood), MSE (Mean Squared Error), Pseudo R² (Pseudo Coefficient of Determination), R² (Coefficient of Determination), RMSE (Root Mean Squared Error), Sensitivity (Sensitivity), TSS (True Skill Statistic).

### 3.5 Ecosystem types

The most commonly studied ecosystems were high mountain forests (12%) and coastal ecosystems (10%), followed by temperate forest (9%) and urban parks (7%). Most of these studies were carried out in partially or fully protected areas (74%), highlighting the importance of protected areas not only for biodiversity conservation, but also for CES (Figure 8).

**Figure 8.**
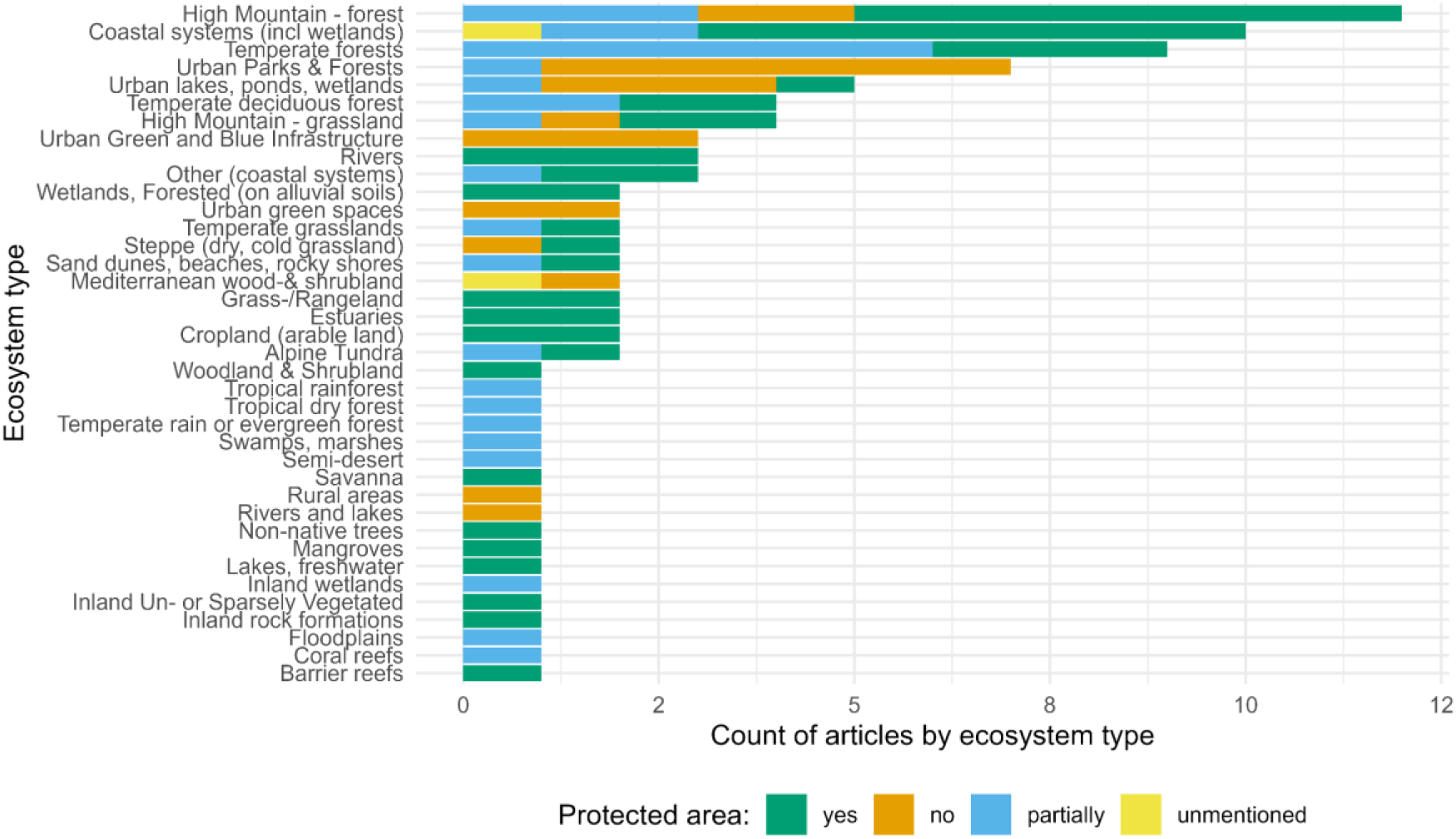
Types of ecosystems studied in the analyzed articles and the proportion of studies in protected areas. Ecosystem types were grouped according to the ecosystem classification provided by The Economics of Ecosystems and Biodiversity (TEEB) (Kumar 2012; Brander et al. 2024).

## 4. Discussion

In this paper, we present a summary of the methods and main variables used in spatial modeling of CES using social media data as a proxy for CES mapping. We found that, in the analyzed papers, a large percentage of CES (28%) are not clearly defined, which generates interoperability problems between publications. The most studied CES are those related to physical and psychological experiences (especially recreation), where the main predictor variables were the presence of natural elements, land use and land cover maps, and topographic variables, although the selection of variables mainly depends on the type of CES evaluated. Several social media data sources were used for information retrieval, with Flickr being by far the most widely used (Manley et al. 2022; Cheng 2023). Although several methods are used to spatially model CES, MaxEnt and Random Forest are the most commonly used.

We found that the most studied CES are those related to physical and psychological experiences (61 %). This is in line with what has been found by other authors (Milcu et al. 2013; Nowak-Olejnik et al. 2022: Márquez et al. 2023; Cheng 2023). However, this emphasis on recreational CES may not only reflect their actual prevalence or importance but may also be influenced by a methodological bias associated with the use of social media data. Social media platforms tend to capture activities that are highly visible, shareable and often associated with leisure and recreation, while potentially overlooking other less tangible CES, such as spiritual connections or knowledge systems. This highlights the need for complementary approaches, as interviews, to ensure a more comprehensive understanding of CES across different contexts and dimensions (Heikinheimo et al. 2017; Vieira et al. 2018; Depietri et al. 2021; Daymond et al. 2023).

We found that in 28% of cases, the author’s categories of CES are oversimplified or incorrectly named since they did not correspond exactly to the precise definition given by the authors of these CES. An adequate and clear categorization and definition is particularly important for the interoperability of ecosystem services data and an adequate interpretation of the results (Bull et al. 2016; Hardisty et al. 2019). There are multiple ways to categorize CES, but it is imperative that first, the categorization adopted match its definition, and second, that we adopt the most comprehensive approach in defining and categorizing them in order to make them easily comparable and standardize them across studies. Although the use of social media data is attractive because it facilitates access to information on a large scale (Sonter et al. 2016; López et al. 2019; Da Mota et al. 2020; Tenkanen et al. 2020), this requires the use of automatic filters to remove as much noise (non-related CES information) as possible. Subsequently, the main challenge is to accurately attribute photos, text or other elements to a specific CES. This may require manual labeling of the type of CES (e.g. Vaz et al. 2020; Fox et al. 2021) or automatic labeling of photographs using machine learning techniques (Cardoso et al. 2022; Huai et al. 2022; Yee and Carrasco 2024).

In the articles analyzed, Flickr was by far the most used platform for accessing geolocated photo data (Wood et al. 2013; Calcagni et al. 2019; Manley et al. 2022; Cheng 2023). Some reasons may be that Flickr offers open access to data through its API and provides geolocated data with an acceptable margin of spatial error (Tallis and Polasky et al. 2009). Other popular platforms such as X-Twitter and Instagram are more restrictive in the use of data and have limitations in providing geolocated information (Ghermandi and Sinclair 2019). On the other hand, other platforms, such as SINA Weibo are very common, especially in China, and offer high-quality data, but are very limited to the Asian continent (Figure 4). Although Flickr is popular among the studies analyzed, it is not the one with the most users worldwide. In fact, the number of users varies widely between platforms: Instagram (2 billion users), X-Twitter (620 million users) and Flickr (60 million users), with the first two having grown the most in recent years (Richards and Friess 2015). As some authors point out, the user profile of each platform also needs to be taken into account, as each platform has a specific focus (Tenkanen et al. 2017; López et al. 2019): they can be mostly used to share everyday activities and experiences (Instagram), thoughts and ideas (X-Twitter), and/or high-quality professional photos (Flickr).

However, it is also important to note that the use of social media data has certain limitations. Poor connectivity in some parts of the world can lead to significant information gaps (Calcagni et al. 2019; Cheng 2023; Pick et al. 2024). In addition, some platforms have restrictive data policies, such as Instagram or Facebook and X - which recently revised their policies on mass data access through their APIs (Ghermandi and Sinclair, 2019) which further complicates data collection. Furthermore, issues such as errors in the geolocation of photos (Zielstra and Hochmair 2013; Longley et al. 2015) require the use of rigorous filtering methods to ensure data accuracy (Fox et al. 2021). These challenges highlight the need for careful consideration and pre-processing when using social media data for spatial analysis.

In relation to predictor variables, we found that the variables used in each study depended mainly on two factors: (1) the type of CES - for example, for a CES related to landscape enjoyment, the accessibility and the presence of attractive natural elements will be very relevant (Table 1); and (2) the environmental characteristics - for example, in a mountainous region, variables such as elevation, slope and aspect might be prioritised as they significantly influence accessibility and aesthetic qualities, whereas in coastal areas, proximity to the coastline, water quality and marine biodiversity might be more relevant. The environmental context not only shapes the physical characteristics of the landscape, but also influences human perceptions and interactions, so it is essential to adapt the choice of variables to the specific setting being studied (Clemente et al. 2019; Hale et al. 2019). For example, water quality may be very important for a recreational CES related to water activities, but not very important for landscape enjoyment. In some cases, the original variables are used (e.g. elevation maps). However, it is more common to use aggregation methods that are applied to the original variables, with the most common distance being. Although Euclidean distance is often used, in some locations, particularly in mountainous areas, distance calculations that take topography into account, such as cost-distance analysis (Salonen et al. 2014; Martínez-López et al. 2019), may be the most accurate.

Regarding the modelling approach used, MaxEnt and Random Forest were by far the most commonly used models to spatially model CES. This is probably because MaxEnt is a well-established and computationally efficient presence-only method that demonstrates strong predictive performance (Sillero et al. 2021). Furthermore, it is particularly well-suited for handling low-presence sample sizes and can accommodate both continuous and categorical predictors. On the other hand, Random Forest, and particularly, down-sampled Random Forest (i.e., RF trained with the same number of presences (1s) and absences (0s)), has been proved to be one of the best modeling approaches for spatial modelling (Valavi et al. 2022). We found that in general most of the studies (88%) used only one type of model, being the use of multiple models less common. It is worth noting that only one study was found to have implemented a model ensemble approach. This is likely due to the fact that model ensembles do not necessarily offer a clear advantage over individually tuned models (Sillero et al. 2021; Valavi et al. 2022). Besides, as Zurell et al. (2020) argued, the choice of models should be based on the type of data (presence/absence or abundance), the amount of data, the predictor variables. For example, Hale 2019) analyzes the most suitable type of model according to the type of data available and the type of CES.

Finally, regarding model performance, the most common metrics used to compare and select models were AUC-ROC, R^2^ and AIC. The AUC-ROC measures the ability of the model to discriminate between classes, with values close to 1 indicating strong performance and values close to 0.5 indicating a model no better than chance. The coefficient of determination (R² and pseudo-R²) assesses the goodness of fit of the model, with higher values indicating greater explanatory power. Meanwhile, the AIC facilitates model comparison by favoring those with a better balance between fit and complexity, with lower values indicating more parsimonious and efficient models. These metrics depend on the model used, for this reason the use of AUC-ROC (typically used to evaluate Maxent models) predominates, but also because it provides a threshold-independent robust assessment of model discrimination ability, particularly for classification problems (Lobo et al. 2008). On the other hand, R² is often employed when evaluating regression models, as it quantifies the proportion of variance explained by the model. However, its use is sometimes criticized in ecological modeling due to potential overfitting and the assumption of linear relationships, which may not always hold in complex environmental systems. Other commonly used metrics in species distribution modelling such as TSS (Allouche et al. 2006) were used in only a few cases. Despite its advantages in balancing omission and commission errors, the limited use of TSS in the reviewed studies may be due to its dependency on a predefined threshold, which can introduce subjectivity in model evaluation (Sillero et al. 2021).

This literature review provides an overview of the models, the social media databases and the variables commonly used to model CES from social media data. This work identifies biases and limitations and aims to support and improve future studies. As general conclusions of the analysis of the articles dealing with this topic, we can suggest some key points for an adequate mapping of CES using social media data. 1) clear categorization and definition of CES, 2) exhaustive filtering of social media data to exclude unrelated content, 3) accurate classification of social media data into a type of CES, 4) careful selection of variables relevant to the type of CES to be evaluated and of the aggregation methods to be used (e.g. distance, density), 5) multi-model analysis, considering techniques for identifying the best models y (Zurell et al. 2020).

## Author contributions

CJN, AE, and DAS worked on the conceptualization. CJN, AE and MRP worked on the process of selecting, screening and extracting information from the articles. CJN and AE worked on formal analysis. NP, RMLL worked on the revision of the CES and their categorisation. JCPG, JML, NP and DAS contributed to significantly improve the drafting and overall coherence of the article. All authors have contributed to the writing – review & editing.

## Acknowledgements

This manuscript was funded by: EarthCul project (PID2020-118041GB-I00 Spanish National Research and Innovation Plan 2020). Grant BIOD2022_002, funded by Consejería de Universidad, Investigación e Innovación and Gobierno de España and Unión Europea – NextGenerationEU, from Biorefuges (TED2021-130888B-I00) funded by MCIN/AEI/10.13039/501100011033 and European Union and by the European Union NextGenerationEU/ PRTR). JML was funded by the Plan Propio de Investigación (P9) of the University of Granada. NP was funded by the BioCiTrees project, financially supported by EU’s Horizon 2020 (HORIZON-MSCA-2022-PF-01, Project n° 101105829).

