## Supplementary material for "Spatial modeling of Cultural Ecosystem Services from social media data: Systematic review of operability, opportunities, limitations and ways forward": Figures and Tables

All codes for the generation of figures and the databases used are available at: https://github.com/kampax/SocialMedia-CES-Modeling-Review

Table S1. Table of the 58 articles analyzed with the extracted variables

Table S2. Standardized CES

Table S3. Original search for articles in WOS and Scopus after removing duplicate items

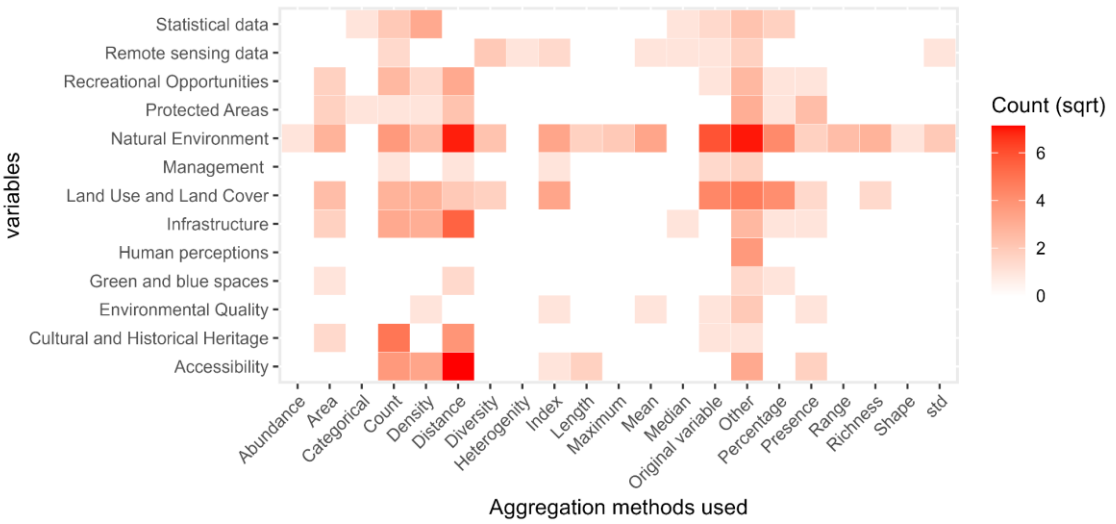


**Figure S1.** Heatmap showing the different aggregation methods used to summarise the variables (x-axis) and the variables used as predictors for the CES modelling, grouped by categories (level 2) (y-axis). The values correspond to the square root (sqrt) of the counts for improved visualisation.

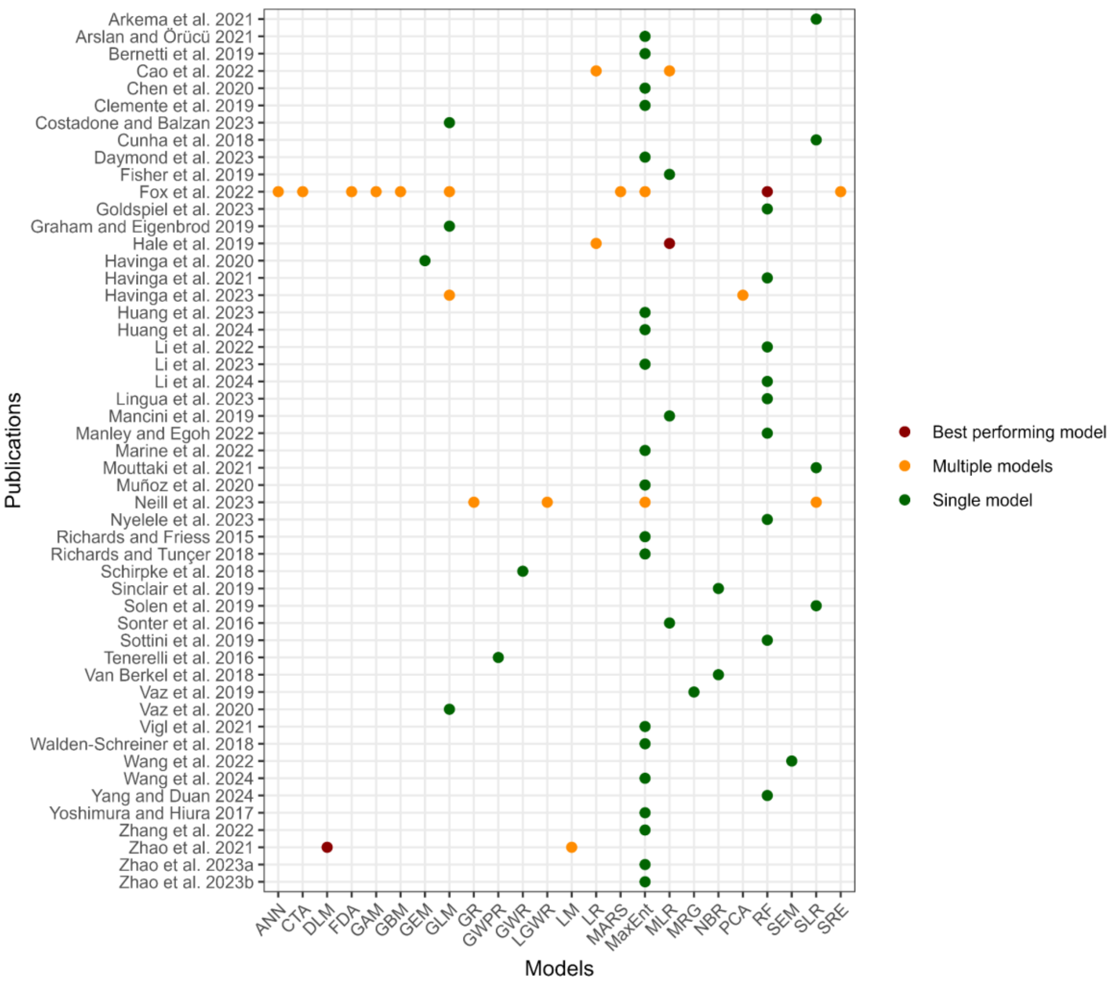


**Figure S2.** Models used in each of the analyzed articles. Articles utilizing a single model are represented in dark green, while those employing more than one model are colored orange. The best performing models are represented in dark red. ANN (Artificial Neural Network), CTA (Classification Tree Analysis), DLM (Dual Logarithmic Model), FDA (Flexible Discriminant Analysis), GAM (Generalized Additive Model), GBM (Generalized Boosting Model), GEM (Global Ecological Model), GLM (Generalized Linear Model), GR (Global Regression), GWR (Geographic Weighted Regression), LGWR (Logistic Geographic Weighted Regression), LM (Logarithmic Model), LR (Logistic Regression), MARS (Multiple Adaptive Regression Splines), MaxEnt (Maximum entropy), MLR (Multiple Linear Regression), NBR (Negative Binomial Regression), RF (Random Forest), SEM (Structural Equations Models), SRE (Surface Range Envelope).

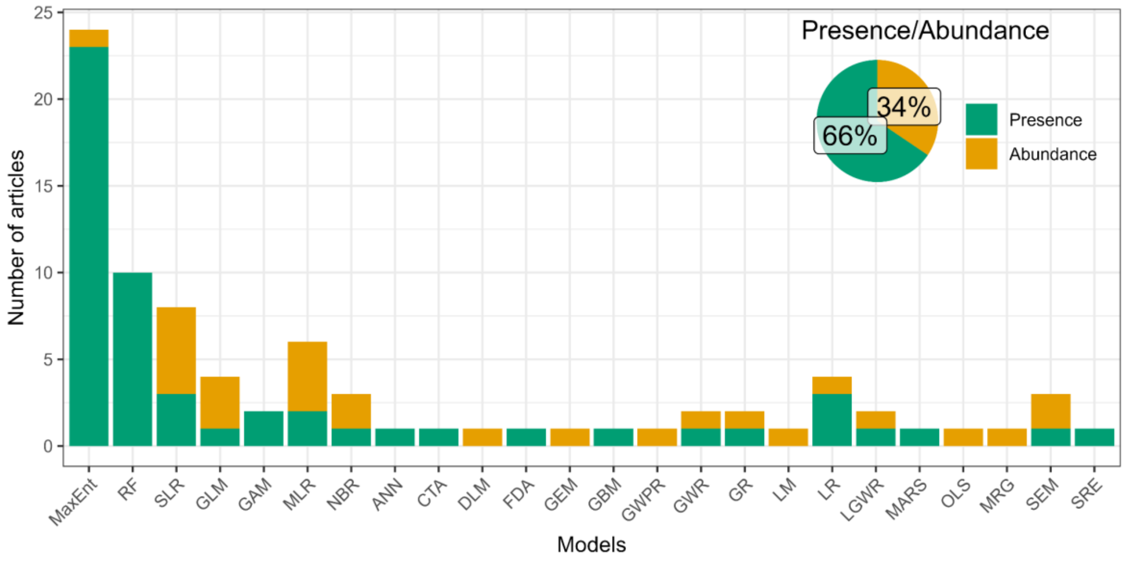


**Figure S3.** Models used and the type of data analyzed. ANN (Artificial Neural Network), CTA (Classification Tree Analysis), DLM (Dual Logarithmic Model), FDA (Flexible Discriminant Analysis), GAM (Generalized Additive Model), GBM (Generalized Boosting Model), GEM (Global Ecological Model), GLM (Generalized Linear Model), GR (Global Regression), GWR (Geographic Weighted Regression), LGWR (Logistic Geographic Weighted Regression), LM (Logarithmic Model), LR (Logistic Regression), MARS (Multiple Adaptive Regression Splines), MaxEnt (Maximum entropy), MLR (Multiple Linear Regression), NBR (Negative Binomial Regression), RF (Random Forest), SEM (Structural Equations Models), SRE (Surface Range Envelope).

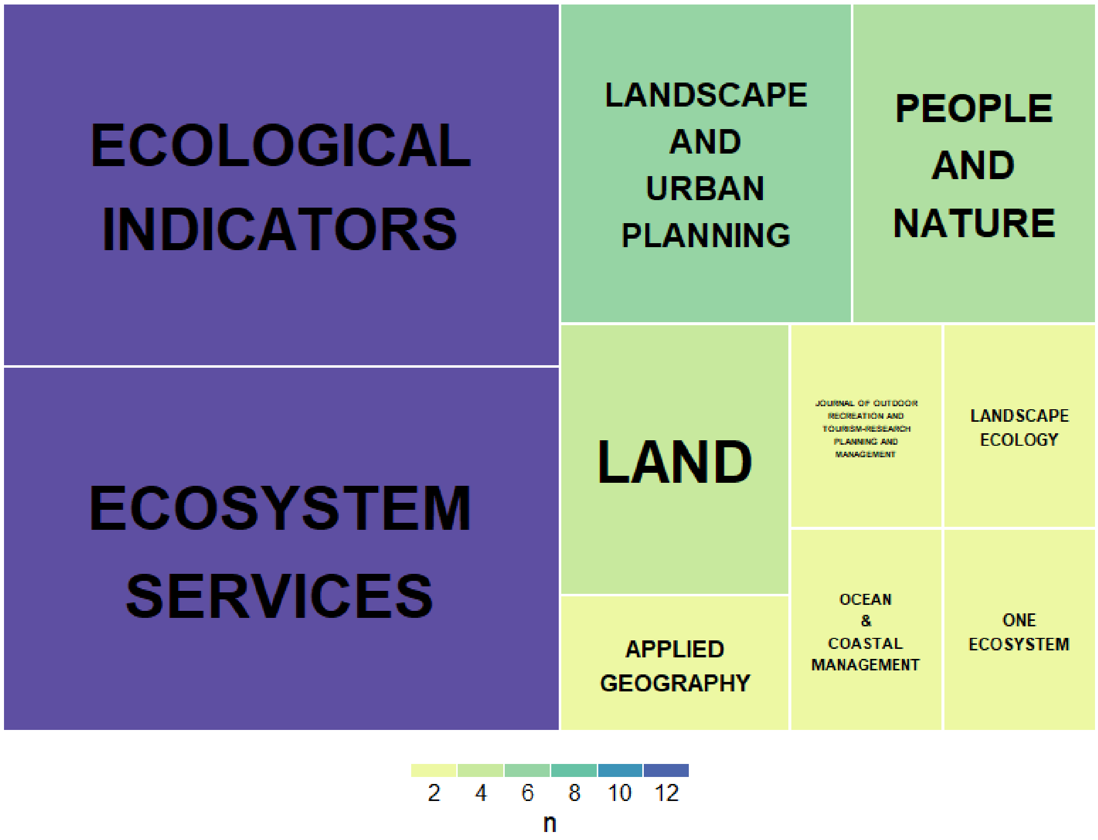


**Figure S4.** Top scientific journals where the analyzed articles were published.

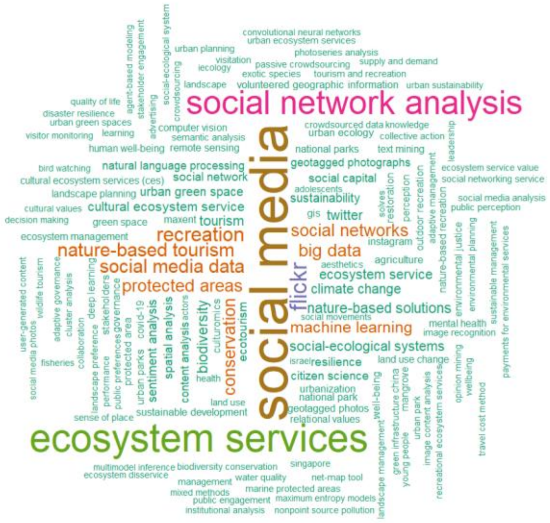


**Figure S5.** Word cloud of the keywords used in the 58 analyzed articles.

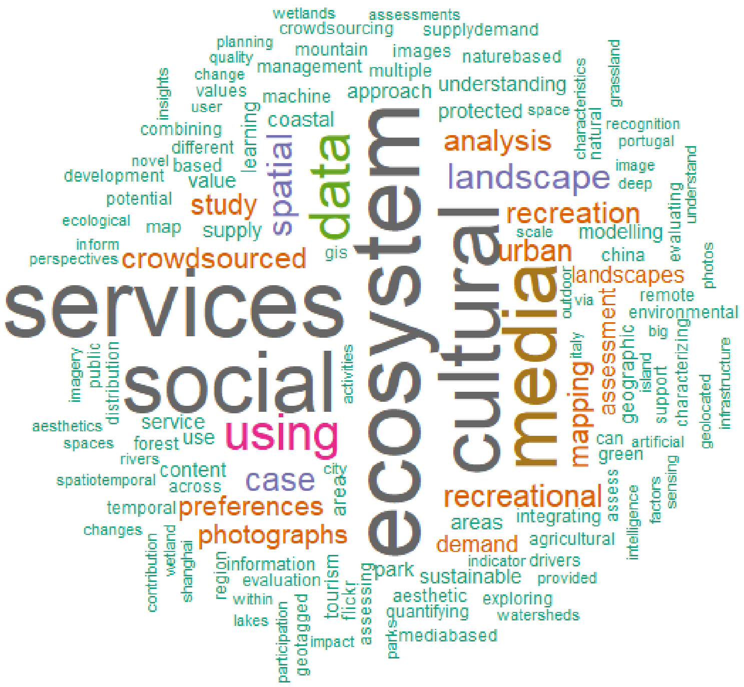


**Figure S6.** Word cloud of the words used in the titles of the 58 analyzed articles.

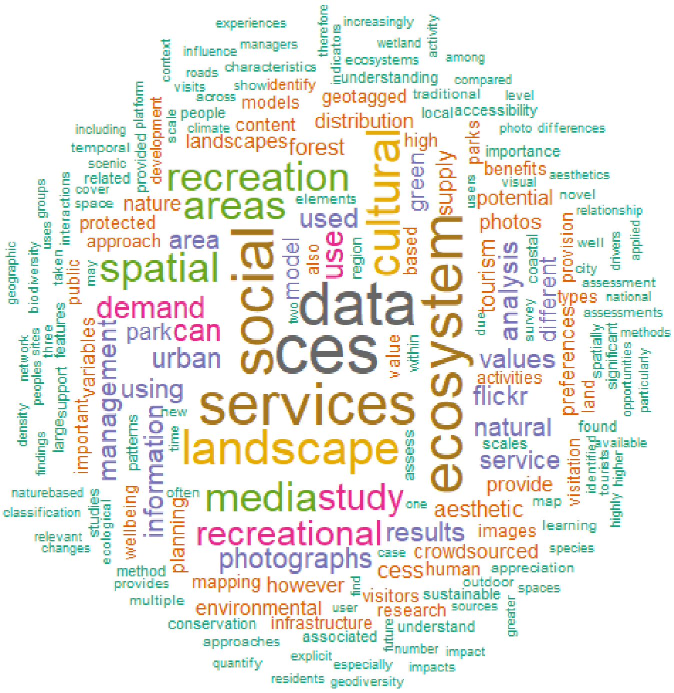


Figure S7. Word cloud of the words used in the abstract of the 58 analyzed articles.

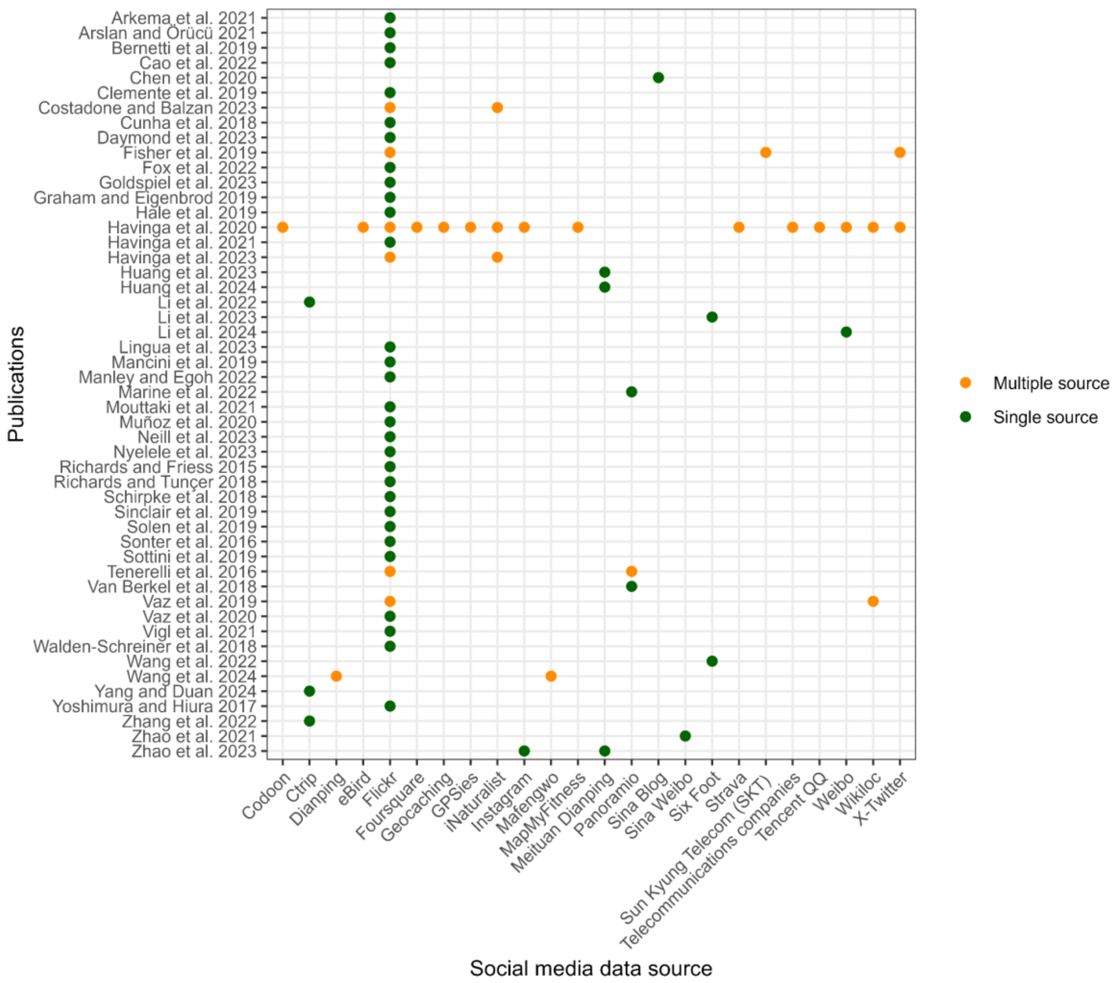


**Figure S8.** Relationship between the articles analysed in this study and the social media data sources used. Orange dots indicate the use of multiple data sources in an article, while green dots represent the use of a single source. Articles are ordered by author and year on the vertical axis.

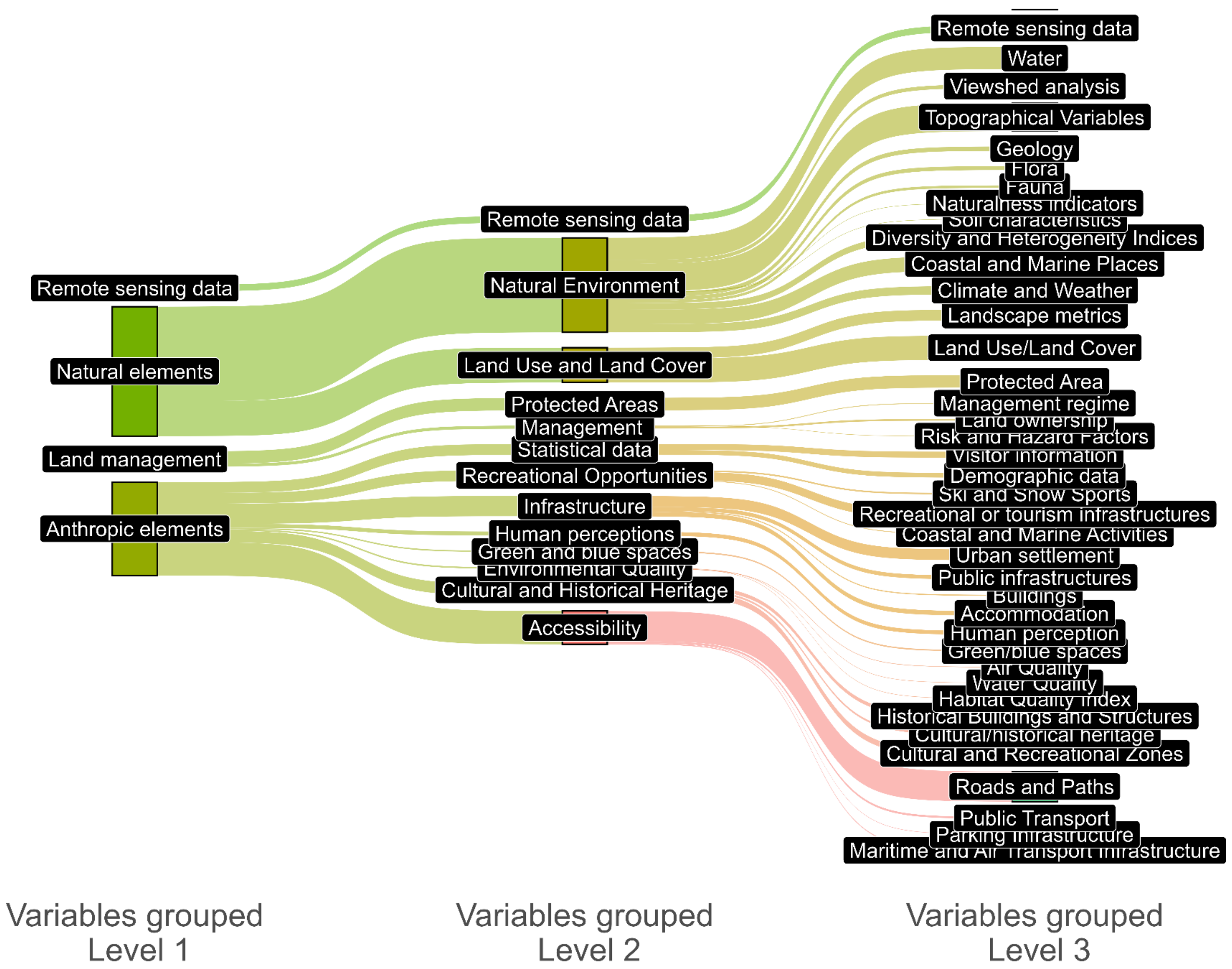


**Figure S9**. Three-level hierarchical classification of variables used as predictors to model CES in the scientific articles analysed. Level 1 is the most general and level 3 the most detailed level of grouping.

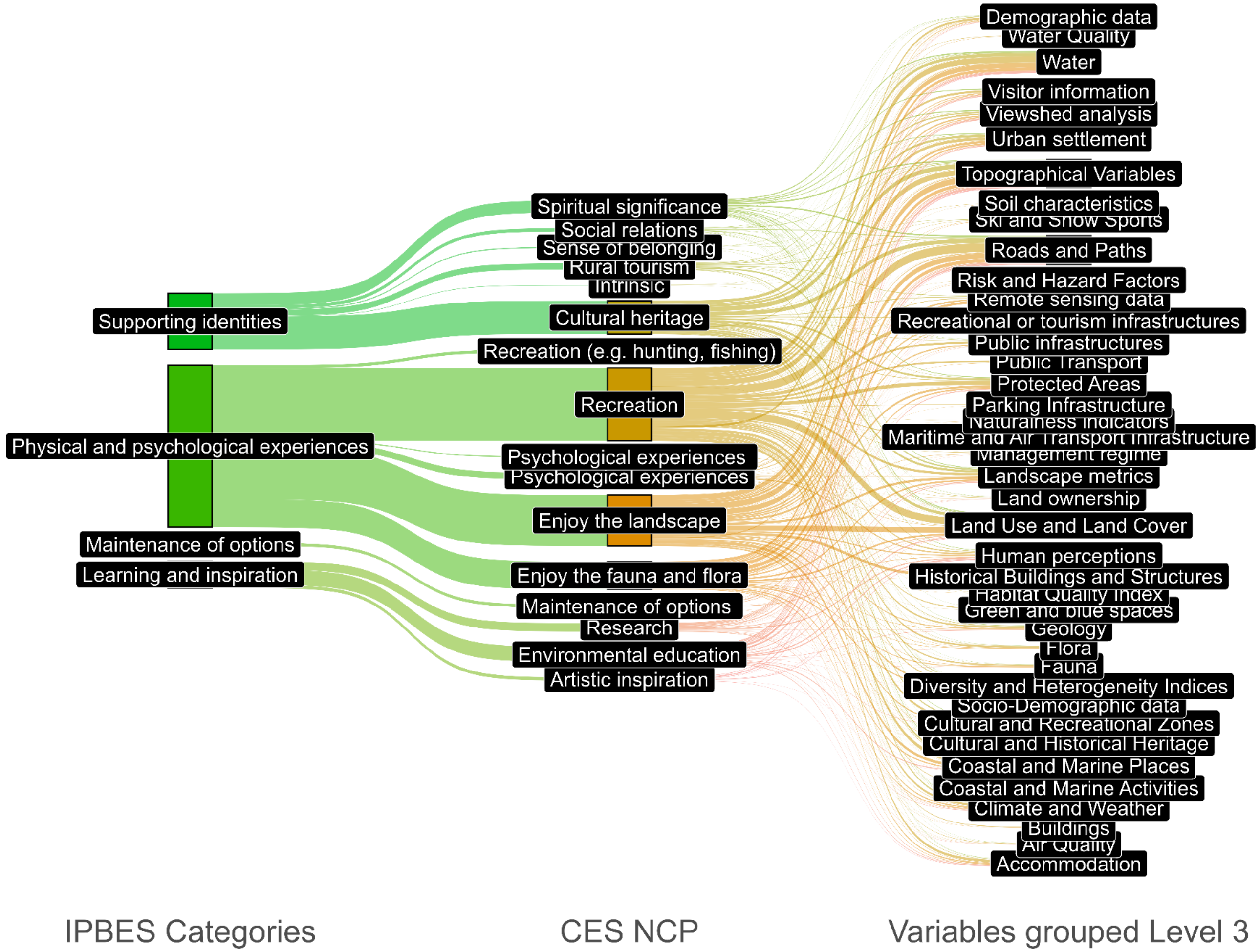


**Figure S10**. Relationship between the variables used as predictors (at the grouping level corresponding to level 3 and the modelled cultural ecosystem services (CES), according to the IPBES (Intergovernmental Science-Policy Platform on Biodiversity and Ecosystem Services) categories and according to our categories.

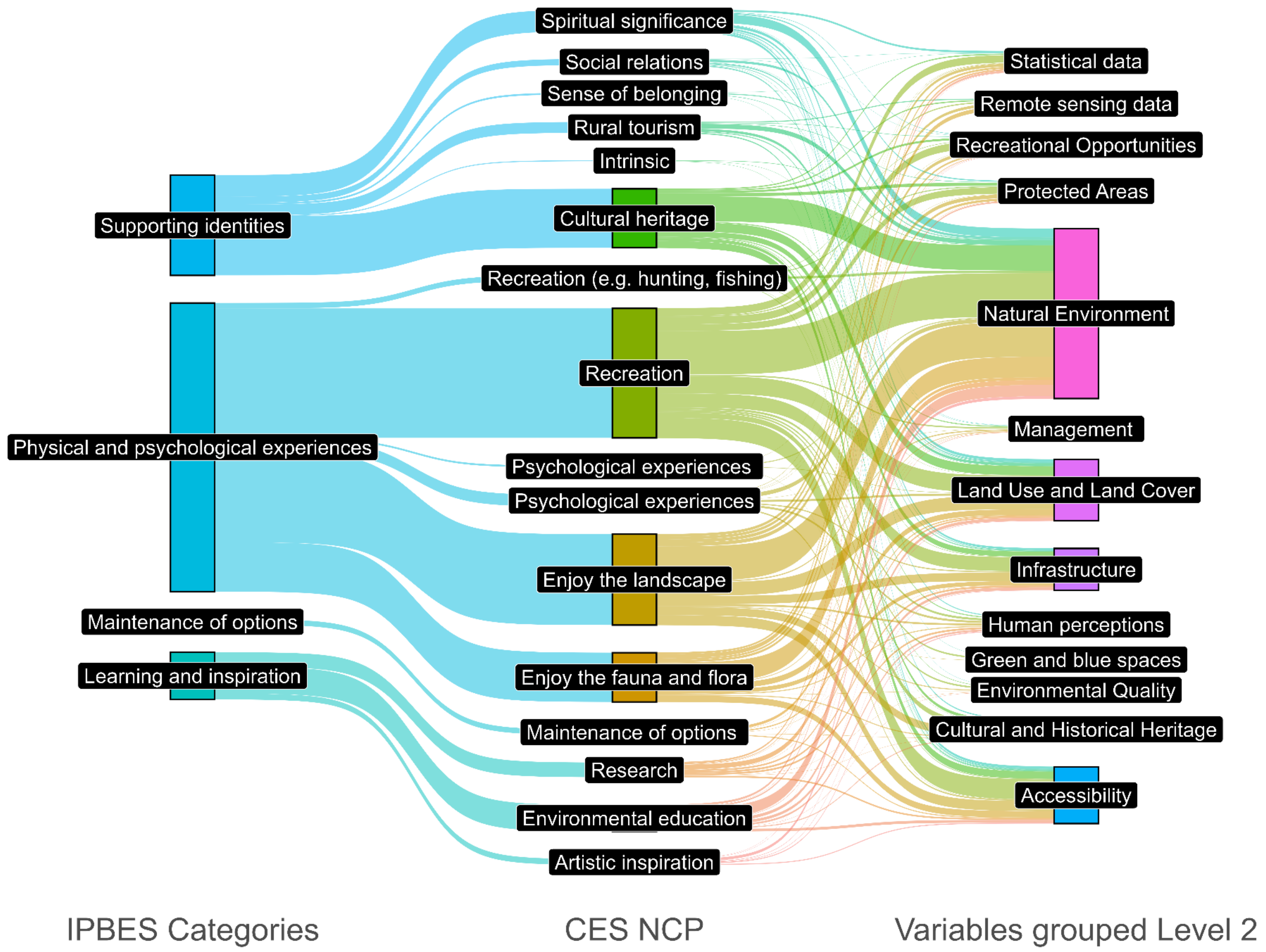


**Figure S11**. Relationship between the variables used as predictors (at the grouping level corresponding to level 2 and the modelled cultural ecosystem services (CES), according to the IPBES categories and according to our categories.
